# Structural Complementarity Maximizes Feasibility and Stability in Microbial Community Coalescence

**DOI:** 10.64898/2026.03.24.713896

**Authors:** Yan Zhu, Bonnie G. Waring, Emma Ransome, Peter Graystock, Bryony E. A. Dignam, Lisa Paruit, Hester J van Schalkwyk, Jie Deng, Thomas Bell, Samraat Pawar

**Affiliations:** Department of Life Sciences, Imperial College London, Ascot, SL5 7PY, United Kingdom; Sir William Dunn School of Pathology, University of Oxford, Oxford, OX1 3RE, United Kingdom

**Keywords:** Microbial community coalescence, Structural complementarity, Feasibility, Dynamical stability, Interaction structure

## Abstract

Microbial communities frequently coalesce through dispersal, disturbance, or deliberate transplantation, yet the dynamical consequences of such coalescence remain poorly understood. Here, we show that coalescence can function as a structural design mechanism to enhance microbial community robustness. Using a mechanistic consumer–resource model in which the balance between competition and metabolic cooperation is explicitly tunable, we quantify how interaction structure shapes both feasibility, namely the environmental domain supporting coexistence, and dynamical stability. Cooperation-dominated communities exhibit greater but more variable feasibility and intrinsic stability than competition-dominated communities. Strikingly, coalescing communities with maximally distinct interaction structures consistently maximizes both feasibility and stability by reducing alignment among interaction vectors and strengthening effective self-regulation in the resulting assemblage. Heterogeneous coalescence balances reduced facilitation, moderated interspecific effects, and stronger self-regulation. These results identify structural complementarity as a general principle for assembling robust microbial ecosystems and provide a theoretical foundation for microbiome engineering strategies that enhance persistence and functional stability.

## 1 Introduction

Microbial communities are rarely isolated systems. In natural ecosystems, they are continuously reshaped by immigration, environmental mixing, and disturbance, with species arriving from surrounding habitats and entire communities coming into contact with one another [Weiss et al., 2023]. Such encounters generate community coalescence events in which two previously independent microbial assemblages mix, reorganizing composition, function, and ecological interactions [Echenique-Subiabre et al., 2025, Amundson et al., 2025, Custer et al., 2024, Liu and Salles, 2024]. Community coalescence also underlies many applied interventions, including probiotic administration, fecal microbiota transplantation, and soil microbiome restoration. Despite its ubiquity, we lack a mechanistic theory linking interaction structure to the robustness of the resulting coalesced ecosystem.

Robust microbial communities must satisfy two fundamental criteria. First, they must be *feasible*: species coexistence must persist across a range of environmental conditions. Second, they must be dynamically *stable*: following perturbation, the community must return to its original state (or equilibrium). Although these properties have been extensively studied in classical coexistence and stability theory, they are rarely quantified explicitly in the context of community coalescence. Most studies focus on which parental community dominates, which taxa persist, or how biomass contributions are redistributed [Diaz-Colunga et al., 2022, Sierocinski et al., 2017, Bresciani et al., 2025]. By contrast, whether coalescence enhances or undermines environmental and dynamical robustness remains poorly understood.

Competitive and cooperative interactions are pervasive in microbial ecosystems and strongly influence coalescence outcomes. Tikhonov [2016] demonstrated that competitive processes such as community-level resource-use efficiency can determine invasion resistance, even in purely competitive systems. Building on this, Lechón-Alonso et al. [2021] showed that cooperative processes (like cross-feeding) can confer advantages during coalescence by enhancing resource-use complementarity. Experimental and theoretical studies further highlight the importance of interaction structure, including competition, facilitation, and species co-selection, in shaping coalesced communities [Diaz-Colunga et al., 2022, Skwara et al., 2023, Albright et al., 2020, Mallon et al., 2018]. However, interaction structure is typically characterized *a posteriori*, inferred from observed networks or random matrices [Tikhonov, 2016, Lechón-Alonso et al., 2021, Serván et al., 2018], rather than treated as a controllable design variable.

As a result, we still lack a systematic understanding of how variation in the competition–cooperation balance of parental communities shapes the feasibility and stability of coalesced communities. In particular, it remains unclear whether robustness is maximized when parental communities are structurally similar or when they differ substantially in their interaction structures. Theory suggests that mixtures of antagonistic and facilitative interactions can enhance diversity and stability relative to single-mode systems [Mougi and Kondoh, 2012]. In microbial contexts, strong competition can stabilize communities by imposing negative feedback, whereas excessive facilitation can destabilize dynamics through reinforcing positive loops [Coyte et al., 2015]. Natural communities typically contain both interaction types simultaneously [Hoek et al., 2016, D’Souza et al., 2018], suggesting that structural heterogeneity may play a central role in ecological robustness.

Here, we treat interaction structure as a continuously tunable design parameter. Using a general mechanistic mathematical model, we explicitly control the balance between competition and metabolic cooperation and systematically examine how the structure of parental interactions influences the robustness of the coalesced community. Rather than focusing solely on taxonomic dominance, we quantify both the extent of environmental feasibility and the dynamical stability of coalesced systems across a continuous spectrum of interaction structures.

We hypothesize that structural heterogeneity between parental communities enhances the robustness of the coalesced ecosystem. Specifically, we test whether combining competitive and cooperative structures embeds stabilizing negative feedback into facilitative networks while expanding the environmental domain that supports coexistence. By explicitly linking interaction design to both feasibility and stability, our framework identifies general principles governing microbial community coalescence and provides a theoretical foundation for engineering robust microbiomes.

## 2 Results

We generated parental communities with prescribed interaction structures by varying the cooperation-to–competition balance parameter *b* (Fig. 1). This parameter simultaneously controls the uptake matrix (**U**) and leakage matrix (**L**) in the MiCRM. As *b*→0, communities are cooperation-dominated, characterized by modular resource specialization and high metabolic leakage. As *b*→ 1, communities are competition-dominated, with overlapping resource preferences and reduced leakage.

**Figure 1.**
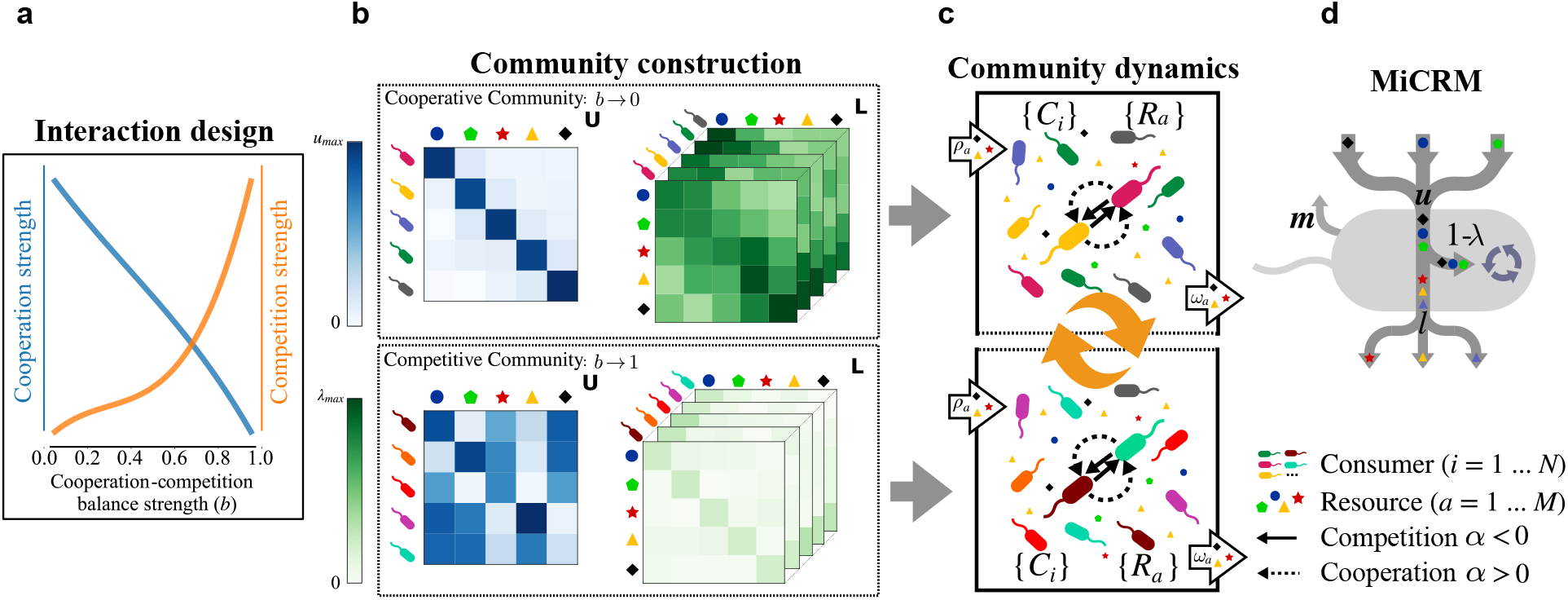
Framework for structurally controlled community coalescence. (**a**) A balance parameter *b* continuously tunes interaction structure from cooperation-dominated (*b*→0) to competition-dominated (*b*→1) regimes. (**b**) *b* modulates uptake (**U**) and leakage (**L**): low *b* weakens competition and enhances cross-feeding, whereas high *b* strengthens competition and reduces cross-feeding. The two parental communities contain distinct consumer species, which are distinguished by color, while the shared resource types are distinguished by geometric symbols. (**c**) Parental communities are simulated to a stable state and coalesced pairwise across (*b*_1_, *b*_2_) to evaluate coalesced-community robustness. Consumer and resource symbols used throughout the schematic are shown below panel **d.** (**d**) In the MiCRM, consumed resources are partitioned into a retained fraction (1 - *λ*), which is processed intracellularly and supports biomass production and growth (blue circular arrows), and a leaked fraction (*l*) that generates byproducts (grey bottom arrows) mediating indirect species interactions (details in section 4.1).

For each value of *b*, we simulated parental communities to a stable state (i.e., equilibrium) and then coalesced all combinations (*b*_1_, *b*_2_) to form coalesced communities. We quantified dynamical stability from the dominant eigenvalue of the Jacobian matrix and environmental feasibility using a derived generalized Lotka–Volterra (gLV) representation. The gLV approximation closely reproduces MiCRM equilibrium dynamics across interaction extremes (Fig. 2a), validating its use for feasibility analysis.

**Figure 2.**
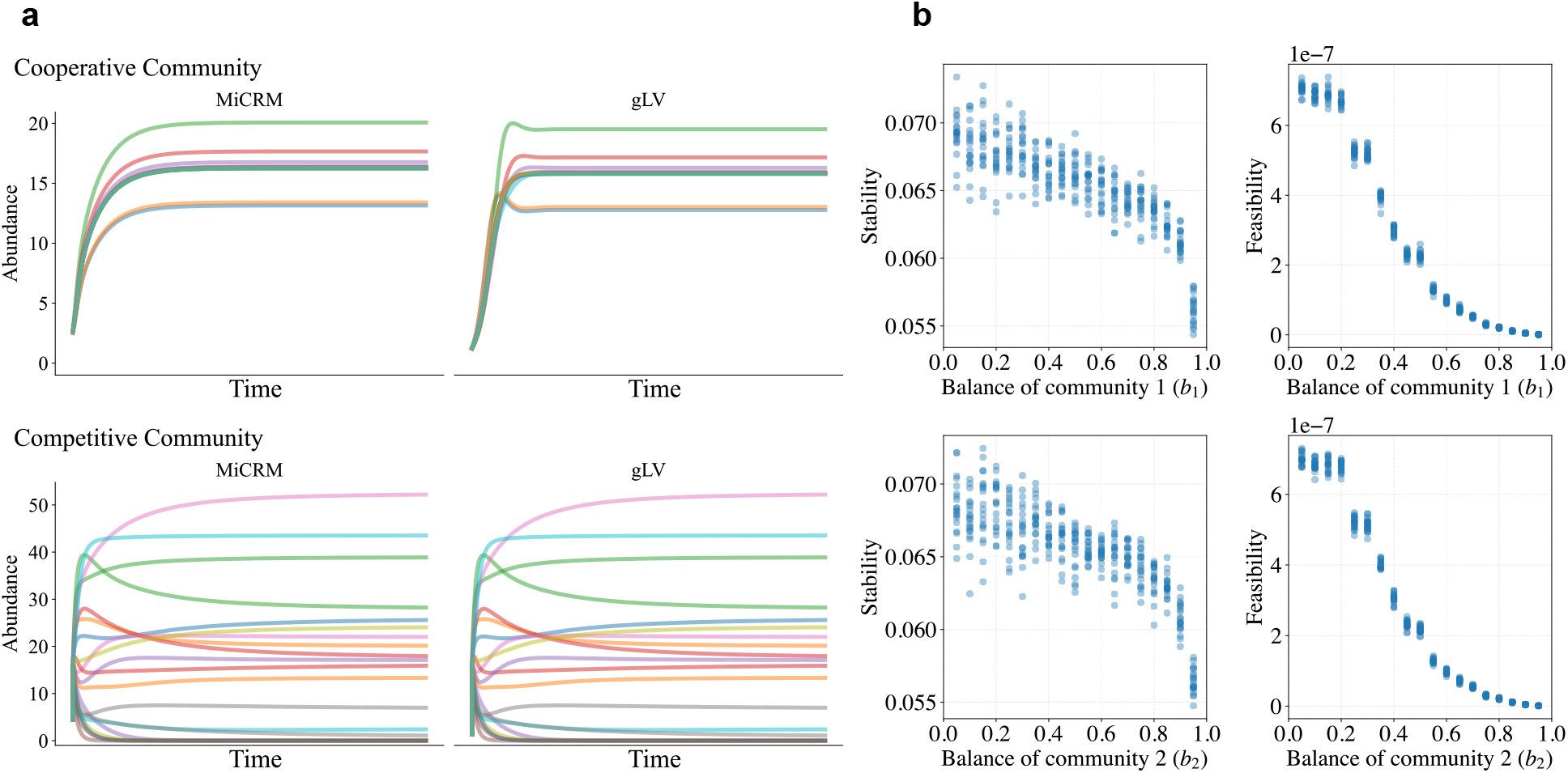
Parental community interaction structure shapes robustness. (**a**) Biomass trajectories (colored curves) from the full MiCRM and the derived gLV model under strongly cooperative (*b* = 0.001) and strongly competitive (*b* = 0.999) regimes converge at equilibrium. (**b**) Stability (the magnitude of the dominant Jacobian eigenvalue) and feasibility (the size of the environmental coexistence domain) decline as the interaction structure shifts from cooperation toward competition.

### 2.1 Interaction structure shapes parental robustness

We first quantified how interaction structure alone affects robustness. Both dynamical stability and environmental feasibility decline as communities shift from cooperation-dominated to competition-dominated regimes (Fig. 2b). Cooperation-dominated systems (*b*→ 0) exhibit greater stability and substantially larger feasibility domains, indicating greater resilience to perturbation and broader environmental tolerance. In contrast, competition-dominated communities (*b*→ 1) display reduced stability and sharply contracted feasibility domains.

Although stability and feasibility covary along the interaction gradient, they decline with different curvature, indicating a trade-o” in how structure constrains environmental *versus* dynamical robustness. Cooperation-dominated communities also exhibit greater variability in robustness across replicates, whereas competition-dominated systems are more compressed and predictable. Under low resource supply (Appendix A), the stability ranking reverses, with competition enhancing stability, while feasibility remains largely unchanged. These results show how interaction structure constrains environmental and dynamical robustness prior to coalescence.

### 2.2 Structural heterogeneity maximizes coalesced robustness

We next examined how robustness may be influenced by structural similarity between parental communities. For each (*b*_1_, *b*_2_) pair, we quantified the feasibility and stability of the coalesced community (Fig. 3).

**Figure 3.**
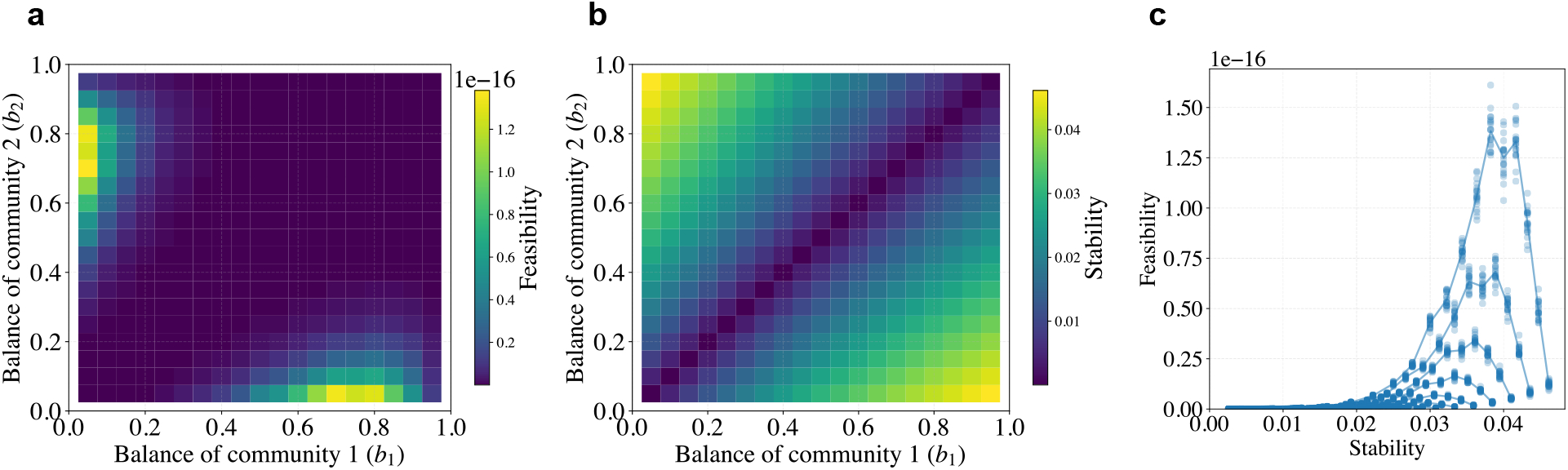
Structural heterogeneity of parental communities maximizes robustness of the coalesced community. (**a**) Environmental feasibility of the coalesced community across parental parameter pairs (*b*_1_, *b*_2_). (**b**) Dynamical stability across (*b*_1_, *b*_2_). Robustness is minimized along the diagonal (structural similarity) and maximized o”-diagonal (structural heterogeneity). (**c**) Relationship between feasibility and stability across all parameter combinations.

Robustness is minimized along the diagonal (*b*_1≃_*b*_2_), where parental communities share similar interaction structures. Coalescing structurally similar systems, whether both competitive or both cooperative, amplifies shared vulnerabilities and leads to the lowest feasibility and stability.

In contrast, robustness is generally maximized when parental communities differ strongly in interaction structure. O”-diagonal regions, particularly those combining a competition-dominated and a cooperation-dominated parent, exhibit the highest feasibility and stability values (Fig. 3a and b). Although feasibility and stability are not perfectly aligned across parameter space, indicating a trade-o” between them (Fig. 3c), heterogeneous coalescence expands both the environmental coexistence domain and dynamical resilience relative to homogeneous coalescence.

These patterns persist when parental communities share 25% or 50% of their species and when coalescence occurs away from a stable state (Appendix B), demonstrating that structural heterogeneity robustly enhances the stability and feasibility of coalesced communities.

### 2.3 Competitive structures dominate biomass while cooperative structures sustain diversity

To understand how robustness emerges from heterogeneous coalescence, we decomposed the contributions to equilibrium biomass and diversity by parental origin (Fig. 4).

**Figure 4.**
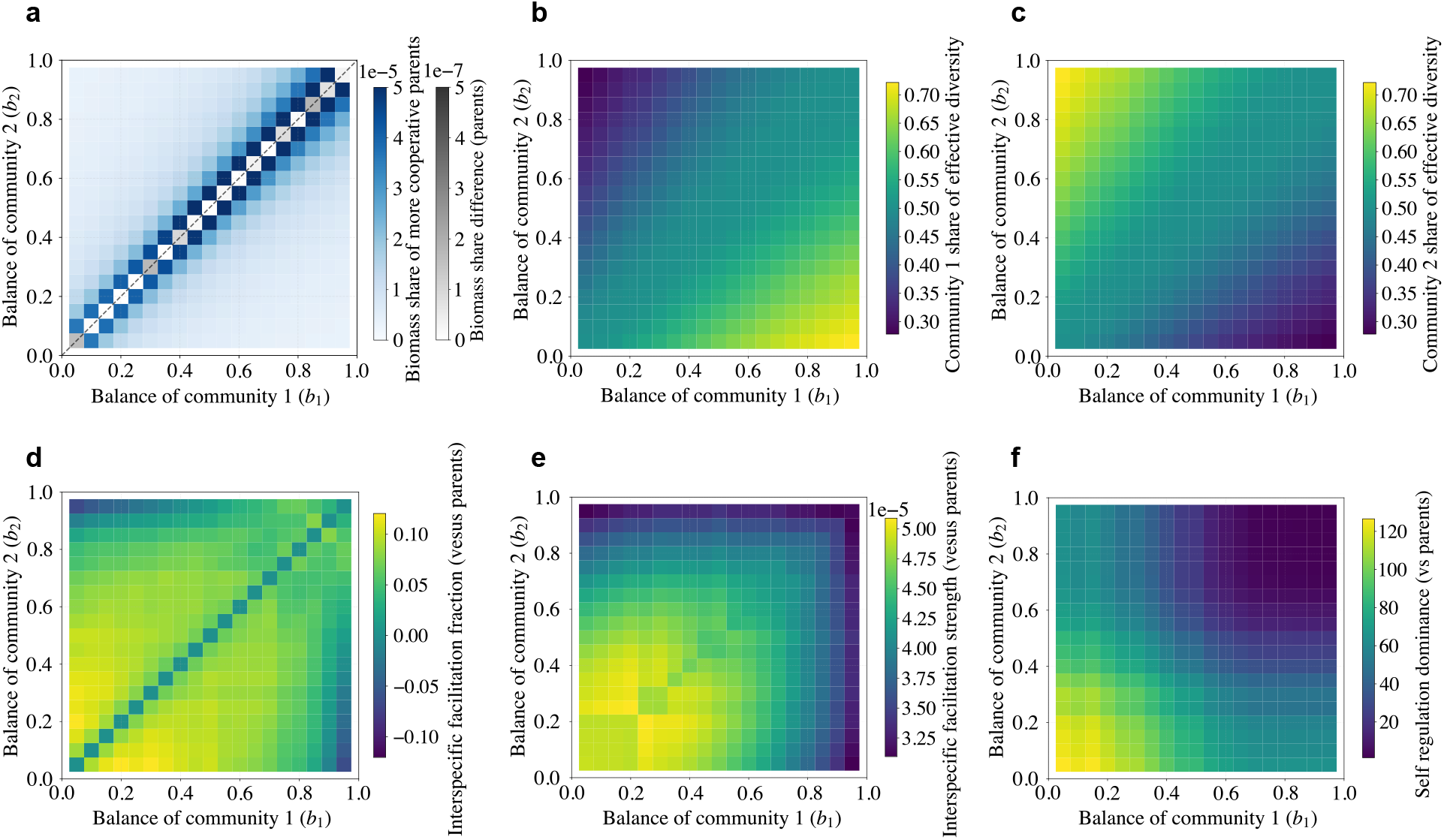
Structural change in the interaction matrix after coalescence. (**a**) Biomass fraction contributed by the more cooperative parent across (*b*_1_, *b*_2_); the more competitive parent contributes the complementary fraction; diagonal values reflect near-equal parental contributions. (**b, c**) Simpson-equivalent effective number of species (defined as 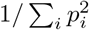) decomposed by parental origin, showing diversity contribution by the two parents. (**d**) Change in fraction of positive interspecific interactions relative to parental mean. (**e**) Change in mean absolute interspecific interaction strength. (**f**) Change in diagonal dominance (self-regulation). Heterogeneity dampens the coalescence-driven amplification of facilitation while moderately enhancing self-regulation, thereby balancing feasibility and stability.

Across parameter space, the more competitive parental community in each pair contributes to a larger part of the total biomass in the coalesced system, even when it is only slightly more competitive than its counterpart (Fig. 4a). Competitive structures therefore provide the dominant biomass backbone of the coalesced community, while cooperative structures expand the space of viable metabolic interactions that support coexistence.

We quantified diversity using the “Simpson-equivalent” effective number of species, defined as 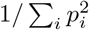, where *p*_*i*_ is the relative abundance of species *i*. This metric converts Simpson’s diversity index into the number of equally abundant species that would generate the same diversity value, providing an interpretable measure of effective diversity. Decomposing this quantity by parental origin (Fig. 4b and c) shows that species from the relatively cooperative parent persist at lower abundance but contribute non-zero effective diversity in the coalesced community. Thus, although more competitive structures dominate biomass, cooperative-origin species expand compositional breadth.

### 2.4 Heterogeneous coalescence restructures interaction structure

Finally, we examined how coalescence alters the interaction matrix itself. Relative to the parental mean, heterogeneous coalescence reduces the prevalence of positive interspecific interactions (Fig. 4d), dampening excessive facilitation. Coalescence increases overall interaction strength (Fig. 4e), but this amplification is constrained by the more competitive parent. Crucially, self-regulation increases across parameter space (Fig. 4f), shifting the community toward greater stability, while heterogeneous coalescence has a more moderate effect on improving self-regulation.

Homogeneous cooperation–cooperation coalescence simultaneously amplifies interaction (Fig. 4d) and its strength (Fig. 4e), increasing fragility. Homogeneous competition–competition coalescence produces minimal structural change (Fig. 4d, e and f). In contrast, heterogeneous coalescence balances interaction amplification with strengthened self-regulation, expanding the feasible environmental domain while enhancing dynamical stability. Yet strong self-regulation can also push equilibrium abundances toward vanishingly small values that are not robust to noise, and weak interspecific coupling can eliminate facilitative buffering, so the feasibility optimum need not coincide with the stability optimum (Fig. 3). Overall, coalescence acts as a structural recombination mechanism that can increase both environmental and dynamical robustness.

## 3 Discussion

Our study identifies structural complementarity as a general feature governing microbial community coalescence. Communities with maximally distinct interaction structures, with one more competitive and the other more cooperative, consistently produce coalesced systems that are both environmentally feasible and dynamically stable. In contrast, coalescing structurally similar communities amplifies shared vulnerabilities and reduces robustness. Coalescence, therefore, acts not merely as a mixing process but as a structural recombination mechanism that can greatly influence community resilience.

Our results distinguish two dimensions of robustness. Feasibility captures robustness to environmental variation, quantified as the volume of intrinsic growth-rate space that supports coexistence. Stability quantifies robustness to dynamical perturbations, as measured by the spectral properties of the Jacobian matrix. Cooperation-dominated parental communities exhibit greater but more variable feasibility domains and stability under sufficient resource availability, whereas competition-dominated communities are more predictable in their robustness but persist only under narrower environmental conditions. These intrinsic constraints define the robustness of parental communities prior to coalescence.

Mechanistically, the structure of the interaction network shapes robustness. Cooperative communities partition resources among species more distinctly, which reduces niche overlap and allows coexistence across a broader range of environmental conditions. In competition-dominated communities, species rely on more similar resources, so niche overlap increases and coexistence becomes restricted to a narrower environmental range, consistent with limiting similarity theory [Levine and HilleRisLambers, 2009, Chesson, 2000]. Stability follows related but distinct structural logic. Strong self-regulation, in which self-limitation terms outweigh interspecific interaction terms, promotes local stability by causing perturbations to decay more rapidly [Allesina and Tang, 2012]. Cooperation increases specialization through metabolic division of labor, but can also amplify interspecific feedback through cross-feeding; competition suppresses facilitation, but increases niche overlap. The observed changes in stability and feasibility along the competition–cooperation gradient therefore emerge from the tension between these opposing structural effects.

Resource supply further clarifies this distinction. Feasibility depends only on intrinsic interaction structure and remains invariant to external input, whereas dynamical stability responds to changes in growth mode and resource availability. Under low-resource conditions, increased reliance on cross-feeding can destabilize cooperative systems, consistent with previous findings [Coyte et al., 2015, Gibbs et al., 2022]. Under abundant supply, specialization reduces indirect competition and enhances stability. Thus, environmental context modifies dynamical resilience but does not alter the geometric feasibility constraints imposed by interaction structure.

The most striking result emerges from structural asymmetry. When two parental communities share a similar interaction topology, their Jacobian spectra exhibit comparable structure, and coalescence reinforces existing unstable modes. In contrast, heterogeneous coalescence embeds stabilizing negative feedback from competitive structures into facilitative networks while reducing alignment among interaction vectors. This dual effect shifts eigenvalues toward stability and simultaneously widens the feasibility domain. In other words, structural heterogeneity makes coalesced communities more resilient to perturbation and able to persist across a broader range of environmental conditions.

Biomass and diversity partitioning further reveal a division of structural roles. The more competitive parent supplies the dominant biomass backbone of the coalesced community, whereas the cooperative par-ent contributes low-abundance but functionally complementary species that expand effective diversity. This asymmetric contribution suggests that competitive structures stabilize abundances, while cooperative structures broaden coexistence. Robustness, therefore, emerges not from a single interaction mode but from their structured combination.

These findings align with empirical observations. Natural and synthetic microbiomes frequently contain mixtures of competitive inhibition and cross-feeding facilitation rather than extreme regimes [Bakkeren et al., 2025]. Systems combining antagonistic and facilitative interactions often exhibit enhanced coexistence and resilience relative to homogeneous networks [Weiss et al., 2022, Niehaus et al., 2019]. Our results provide a mechanistic explanation for this pattern: structural complementarity expands environmental tolerance while embedding stabilizing feedback.

Beyond explaining coalescence outcomes, this framework suggests design principles for microbiome engineering. Introducing consortia with complementary interaction structures, rather than structurally similar communities, may enhance persistence, invasion resistance, and functional stability. Competitive modules can constrain runaway facilitation and stabilize biomass, whereas cooperative ones can expand metabolic complementarity and environmental tolerance. Coalescence thus becomes a controllable architectural tool for assembling robust microbial ecosystems.

Overall, our results demonstrate that the robustness of coalesced communities depends not only on the dominance of cooperation or competition, but also on the heterogeneity of their interaction structures. By linking feasibility geometry and dynamical stability to explicitly tunable structure, we provide a general theoretical foundation for understanding, predicting, and engineering microbial community resilience.

## 4 Methods

### 4.1 The Model

We use the general MiCRM, which for a community of *N* species and *M* resource types reads:

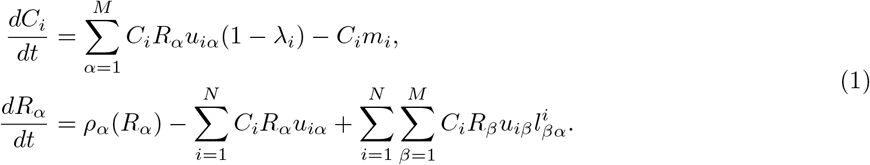

Here, *C*_*i*_(*i* = 1,. .., *N*) denote the biomass abundance of the *i*th species (consumer) and *R*_*α*_(*α* = 1,. .., *M*) the abundance of the *α*th resource type. We define the external resource input function as *ρ*_*α*_(*R*_*α*_) =*ρ*_*α*_ − *R*_*α*_ *ω*_*α*_, where *ρ*_*α*_ is the constant external supply rate of resource *ρ*, and *ω* _*α*_ is the removal rate due to dilution, outflow, or decay. All resources are externally supplied at constant rates *ρ*_*α*_ and are removed from the system through outflow or decay at rates *ω* _*α*_. The growth of species *i* is governed by its total resource uptake and maintenance loss rate *m*_*i*_. The parameter *α*_*i*_ denotes the leakage fraction of species *i*, such that only a fraction (1 − *α*_*i*_) of the assimilated resource contributes to species growth, while the remaining fraction is released as metabolic byproducts. Resource uptake is determined by a preference vector *u*_*iα*_, specifying the affinity of species *i* for each resource *ρ*. Consumed resources are either released back into the same resource type (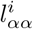) or partially transformed into metabolic byproducts (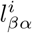), which are released back into the environment. For each species *i* and consumed resource *β*, the byproduct allocation coefficients satisfy 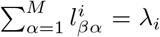 so that *α*_*i*_ sets the total leaked fraction and 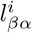 specifies its partitioning among byproduct types. Leakage efficiencies are constrained at both the species and resource levels. The MiCRM is general in that it provides a unified, minimal framework that flexibly captures variation in resource competition and cross-feeding, as well as in energy-use traits and resource-supply types. Stability analysis of the MiCRM is described in Appendix D.

### 4.2 Controlling the cooperation-competition balance

We construct pairs of communities with different and identical interaction structures by controlling a cooperation-competition balance strength parameter (*b* ∈ [0, 1]). This parameter is based on the obser-vation that competitive communities exhibit high niche overlap and reduced resource partitioning, whereas cooperative ones are characterized by modular cross-feeding networks and enhanced complementarity in resource uptake. Therefore, the parameter *b* tunes both the modular structure and the entrywise values of the uptake matrix **U** and the leakage matrix **L**. Specifically, we define the number of consumer-resource modules as:

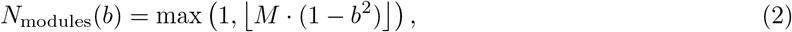

such that cooperative communities (*b* → 0) exhibit modular structure, while competitive ones (*b* → 1) collapse toward global overlap. Because the number of modules is discrete, the ⌊ · ⌋ operation can introduce stepwise jumps as *b* varies. We therefore use a squared dependence on *b* to smooth the parameter sweep by mitigating rounding-induced staircase effects.

The specialization ratio, which governs the degree of resource preference heterogeneity among consumers, is assigned as:

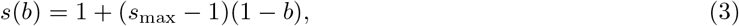

ensuring that low-balance strength yields high specialization (low niche overlap), whereas *b*→ 1 recovers a uniform uptake rate.

Finally, the per-species leakage fraction (*α*) is interpolated linearly:

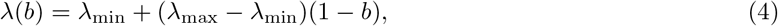

capturing the tendency for cooperative systems to recycle more metabolic byproducts. Therefore, as shown in Fig. 1, varying *b* continuously generates a spectrum of interaction structures from cooperative to competitive: as *b*→ 0, **U** becomes increasingly diagonally dominant so that species partition resources with minimal overlap, and **L** increases to promote stronger leakage and conversion into other resource types; as *b*→ 1, **U** becomes progressively more random, increasing niche overlap, while **L** decreases, corresponding to weaker leakage. The role of the cooperation-competition balance *b* is discussed further in Appendix E.

### 4.3 Effective species interactions

To capture the intrinsic dynamics of microbial communities, we require a model that explicitly describes interspecific interactions. In the MiCRM, interactions are mediated implicitly through shared resources and metabolic byproducts, which limits analytical exploration in high-dimensional environmental space. We therefore coarse-grain the MiCRM into a species-only gLV model. The resulting gLV formulation takes the following form:

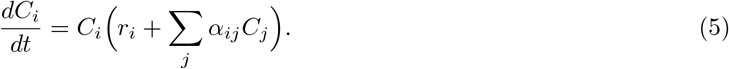

The two key parameters in the gLV model, interaction coefficient matrix (**A** = [*α*_*ij*_]) and intrinsic growth rate (*r*_*i*_), are given by the following equations:

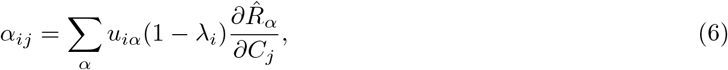

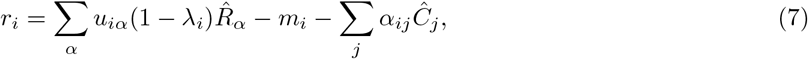

where 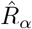 and *Ĉ*_*j*_ denote the equilibrium abundances of resource *α* and species *j*. The detailed derivation is provided in Appendix F.

### 4.4 Coalescence process

First, the two parental communities were simulated independently under the same resource conditions until each reached equilibrium; they differed only in species composition. To simplify the detection of equilibrium, we terminated the simulation when the maximum absolute rate of change across all species fell below a small threshold (1.0 × 10^*→*5^). We then implemented coalescence by combining the two species pools into a single pool that shares the same *M* resource types. Specifically, we concatenated the uptake matrices **U**_1_ and **U**_2_ to form **U**_3_, and concatenated the corresponding leakage tensors **L**_1_ and **L**_2_ to form **L**_3_. We set the initial condition of the coalesced community (community 3) by concatenating the parental equilibrium point, 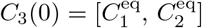. By contrast, the resource state was reset to the same initial resource supply conditions used for communities 1 and 2, with identical inflow and outflow rates. Finally, we numerically integrated community 3 to its equilibrium under the same MiCRM dynamics to investigate how patterns of parental communities influenced the final composition of the coalesced community. Figure 1 provides an overview of the research framework. We assessed the stability of the three communities (two parental communities and one coalesced community) at their equilibrium states to evaluate how the system responds to perturbations.

### 4.5 Coexistence theory

Under the assumptions of coexistence theory, the interaction matrix **A** is treated as an intrinsic property of the community and is therefore invariant to environmental change, whereas the environment acts directly on the intrinsic growth vector **r** [Long et al., 2024]. We represented environmental uncertainty as a parameter space Θ ⊂ ℝ^*N*^ for **r**. In the absence of prior constraints, we took Θ to be the unit sphere *S*^*N→*1^ with a uniform measure, noting that feasibility is homogeneous in **r** so positive rescalings do not affect qualitative outcomes. When prior knowledge exists, Θ can be restricted to biologically plausible sectors, such as the positive orthant, or deformed into anisotropic sets, such as ellipsoids that weight likely directions more heavily. Given nonsingular **A**, we defined the feasibility region *ℱ* (**A**) = *{***r** ∈ ℝ^*N*^ | −**A**−^1^**r** ≻ **0***}*, feasibility is equivalent to the implied equilibrium **c**^*\**^ = −**A**−^1^**r** lying in the positive orthant, which makes *ℱ* (**A**) a]convex cone intersected with #. We then estimated the probability of coexistence as *p*(**A**) = ℙ_**r**∼ Θ_ [**r** ∈ *ℱ* (**A**) by the Genz–Bretz quasi–Monte Carlo evaluation of the multivariate normal orthant probability, which is effective in high-dimensional systems. This probabilistic measure provides a geometrically intuitive and computationally tractable way to compare ecological feasibility across different communities or environmental contexts. Applying this framework to preand post-coalescence states allows us to quantify how the coalescing of communities reshapes their feasible environmental domain, thereby linking interaction structure to coexistence potential.

## 5 Code availability

All code is available at https://github.com/EcoEngLab/coalescence_robustness.

