## Appendix for "Structural Complementarity Maximizes Feasibility and Stability in Microbial Community Coalescence"

### A Considering Low Resource Supply

We examine the relationship between resource supply and community stability (Fig. A1a). External resource input rate directly influences the stability of both cooperative and competitive communities. Under low-resource conditions, stronger cooperation is associated with lower stability, consistent with previous observations. When external resources are abundant, cooperative communities exhibit higher stability than competitive ones. The model therefore extends earlier conclusions by showing that the effect of cooperation depends on the resource context. For both cooperative and competitive communities, the effect of resource supply on stability is non-monotonic. Stability increases with resource input at low supply levels, but once supply exceeds a threshold, stability declines. Increasing resource input thus enhances stability within one range and suppresses it in another. This pattern indicates a shift in the dominant growth mode. When resources are scarce, species rely primarily on metabolites leaked by others. When resources are abundant, reliance on cross-feeding weakens.

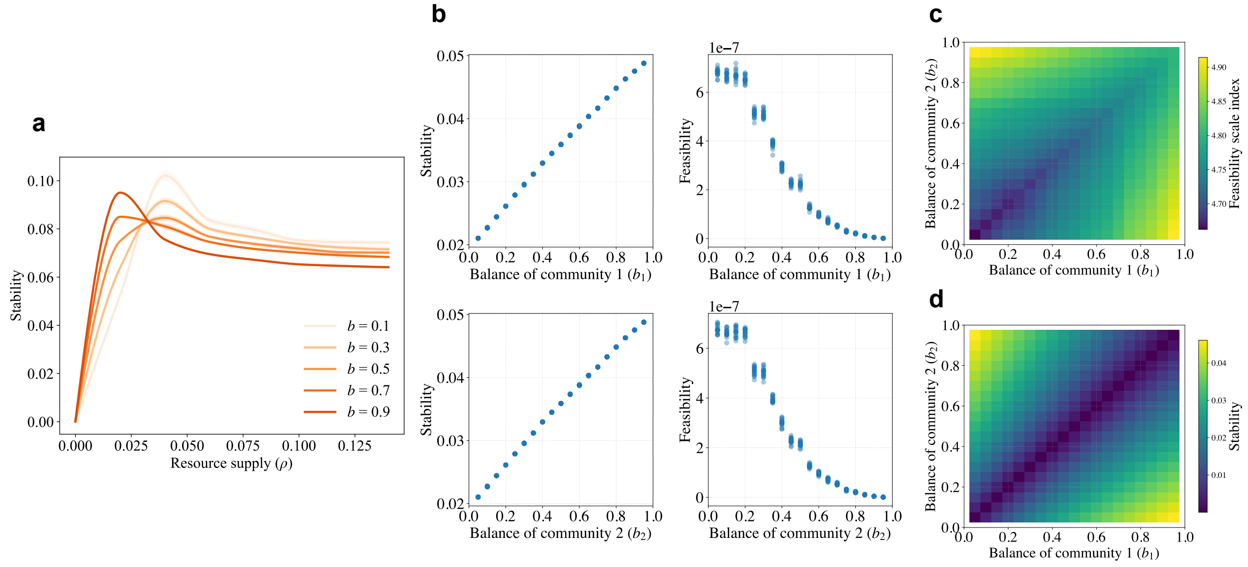


Figure A1: Effects of low resource supply on stability and feasibility. (**a**) External resource input modifies community stability. Lighter orange curves denote more cooperative communities, whereas darker curves denote more competitive ones. Under low-resource conditions, competitive communities are more stable. When resource supply is high, cooperative communities become more stable. Each curve exhibits a peak, and the curves intersect at a turning point, indicating a shift in the dominant growth mode. (**b**) With resource supply fixed at $\rho = 0.01$, variation in $b$ further highlights that competition enhances stability under resource limitation, whereas feasibility remains unaffected by external resource input. (**c**) Heterogeneous coalescence yields higher feasibility than homogeneous coalescence; (**d**) it also maximizes stability of the coalesced community, consistent with the pattern observed under abundant resource supply.

Under low resource supply ($\rho = 0.01$), we analyze how the competition–cooperation balance parameter $b$ affects stability and feasibility (Fig. A1b), as well as coalescence outcomes (Fig. A1c & d). As expected, cooperative communities show lower stability under low resource input, whereas competition enhances stability. Feasibility remains unaffected by the supply of external resources. After coalescence, community dimensionality increases, but limited resource input leads to the exclusion of more species in the coalesced community, making direct estimation of the feasibility-domain probability unreliable. We therefore quantify the size of the feasible domain using a stable spectral/volume proxy based on $\log\det(\mathbf{A}\mathbf{A}^\top)$. This proxy is monotone with the corresponding truncated feasible volume and remains numerically stable across the full parameter scan. We refer to this quantity as the feasibility scale index. Under low external resource supply, both the feasibility scale index and stability reach their maximum under heterogeneous coalescence, consistent with the pattern observed under abundant resource supply.

### B Coalescing Communities with Partially Shared Species

Actually, two parental communities are likely to share a subset of species (i.e., some species occur in both community 1 and community 2), so their uptake matrices $\mathbf{U}$ and leakage tensors $\mathbf{L}$ cannot be concatenated by simple stacking. To examine this scenario, we perform additional coalescing experiments with controlled species overlap. We partition each parental community into shared species and unique species. Shared species are included only once in the coalesced community. Specifically, the coalesced uptake matrix is constructed as $\mathbf{U}_3 = \left[\mathbf{U}_{\text{share}}, \mathbf{U}_{\text{unique},1}, \mathbf{U}_{\text{unique},2}\right]$, and the coalesced leakage tensor $\mathbf{L}_3$ is assembled analogously. For two parental communities each containing $N$ species and sharing a fraction $f \in (0,1)$ of species, the coalesced community contains $N_3 = fN + (1-f)N + (1-f)N = (2-f)N$ species. In our implementation, community 1 is generated under a control parameter $b_1$, community 2 is generated under $b_2$, and overlap is imposed by replacing the first $fN$ species in $(\mathbf{U}_2, \mathbf{L}_2)$ with the corresponding first $fN$ species from $(\mathbf{U}_1, \mathbf{L}_1)$.

In addition, because the equilibrium status of parental communities is typically unknown in empirical settings, we also consider coalescing away from equilibrium. In simulations, this is simplified by coalescing at the initial condition rather than after pre-equilibrating each parental community. When initial abundances are equal across all species in both parents, coalescing at $t = 0$ implies that the initial abundance of each shared species is doubled, reflecting the aggregation of contributions from both communities.

For each $(b_1, b_2)$ pair, we perform 20 stochastic replicates and report replicate-averaged results. Under species overlap, the control parameter $b$ is no longer sufficient to represent the effective strength of competitive–cooperative interactions in parental communities; therefore, we quantify cooperation-competition strength from two complementary perspectives: (i) the mean pairwise cosine similarity among rows of $\mathbf{U}$, and (ii) an effective leakage metric derived from $\mathbf{L}$. In addition, when species are shared, the Gaussian-orthant feasibility probability becomes numerically ill-conditioned and highly variable; small seed-induced perturbations across technical replicates can strongly affect the orthant integral in moderate-to-high dimensions. We therefore apply the feasibility scale index described in Appendix A to quantify changes in feasibility.

Figures A2 and A3 show results for $f = 0.25$ (25% overlap) and $f = 0.50$ (50% overlap), respectively. At both overlap levels, the feasibility scale index and the stability metric exhibit approximately symmetric patterns in $(b_1, b_2)$ space. Specifically, the feasibility scale index is smaller when the two parental communities have more similar uptake matrices, whereas increasing dissimilarity between $\mathbf{U}_1$ and $\mathbf{U}_2$ enlarges the feasibility scale index (see Figs. A2a and A3a). The same qualitative trend is recovered when similarity is assessed via effective leakage. Figures A2c and A3c show the same pattern via effective leakage. Figures A2b and A3b quantify how uptake similarity relates to the stability of the coalesced community, while Figs. A2d and A3d show the corresponding relationship for effective leakage.

Interestingly, comparing Figs. A2 and A3 further shows that increasing species overlap compresses the point cloud toward the diagonal in the $(x, y)$ plane. Intuitively, when overlap is low (including the main-text case of disjoint species sets), the two communities can span a wider range of pairwise dissimilarities, and coalescing outcomes occupy a larger portion of the feasible parameter space. In contrast, when overlap is high (approaching identical communities), coalescence becomes increasingly degenerate, because it induces only limited (or no) structural change relative to either parental community.

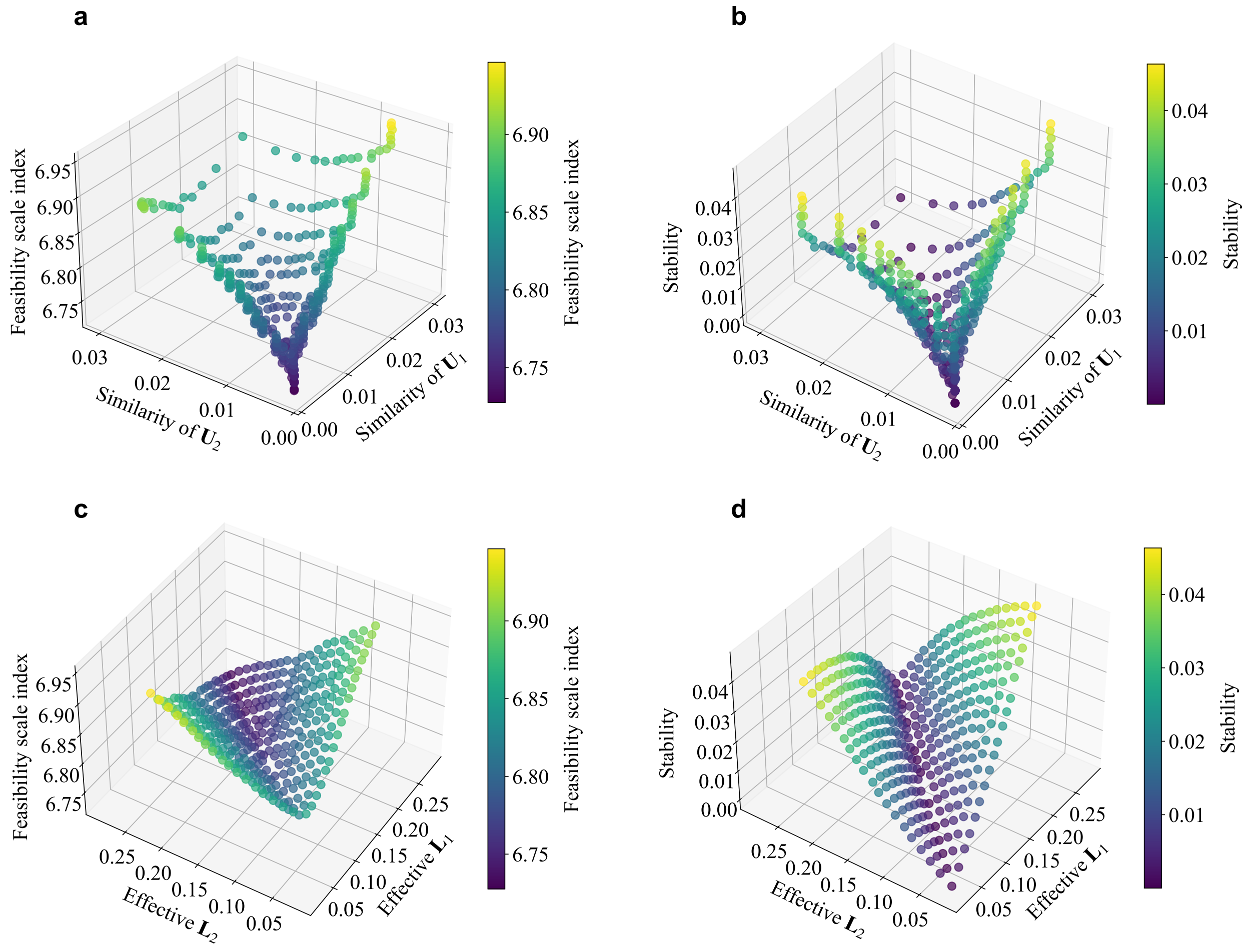


Figure A2: The two parental communities share 25% of the identical species. (**a**) Coalesced-community feasibility scale index as a function of the two parental communities' competitive intensity, quantified by similarity in uptake matrices (higher similarity indicates stronger competition). The surface is approximately symmetric and attains a minimum near equal competitive intensity; feasibility increases with competitive asymmetry. (**b**) Coalesced-community stability versus parental competitive intensity. Stability is lowest near equal competition and increases off the diagonal, consistent with a positive association between stability and feasibility. (**c**) Feasibility scale index versus parental cooperation, quantified by effective leakage (higher values indicate stronger facilitation). Feasibility is minimized near equal cooperation and increases with cooperative asymmetry, mirroring (**a**). (**d**) Stability versus parental cooperation, with a minimum near equal cooperation, paralleling (**b**). Overall, coalescing structurally heterogeneous parental communities increases feasibility and stability, even when the parental communities share 25% of species; the qualitative pattern is consistent whether assessed via competition or cooperation.

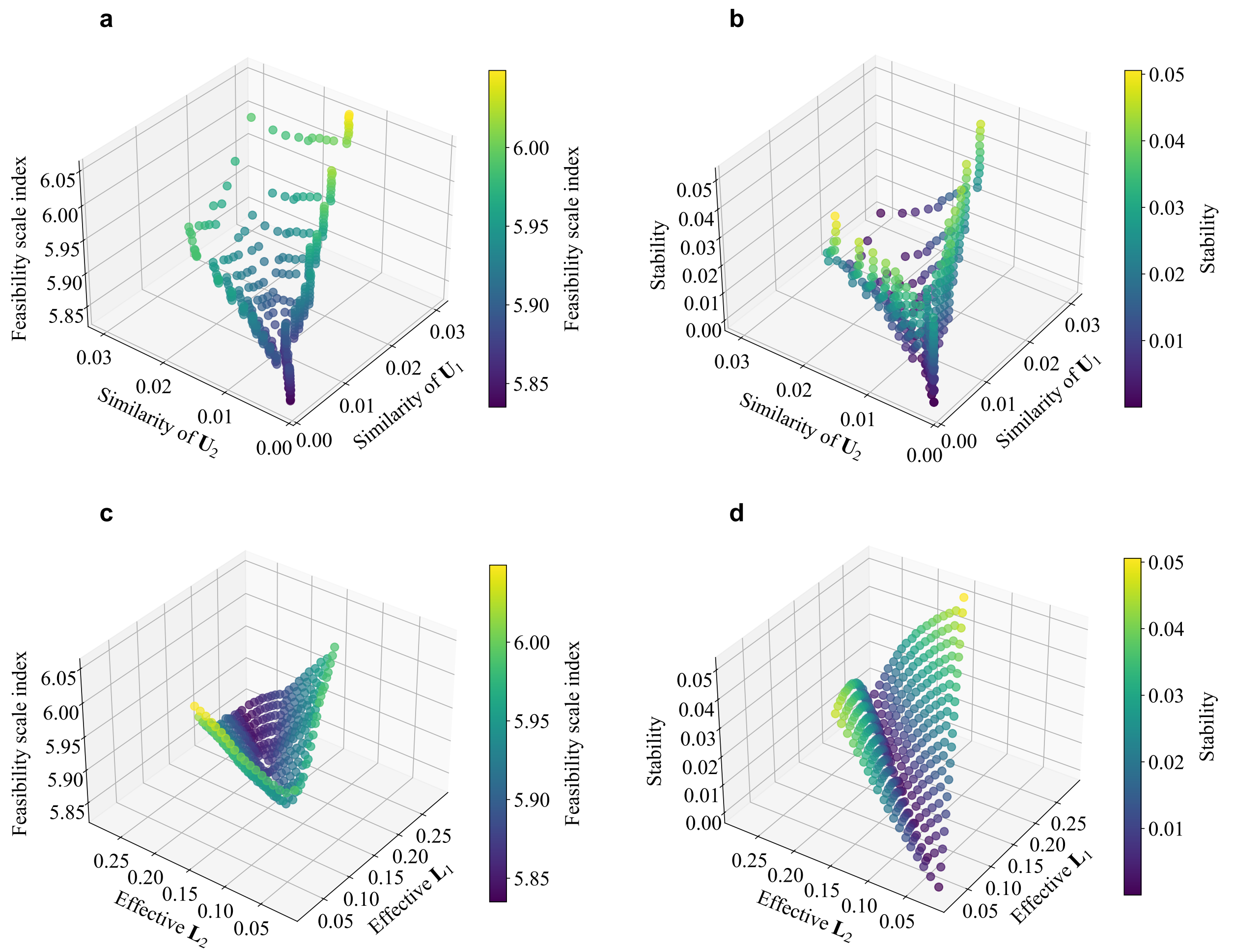


Figure A3: The two parental communities share 50% of the identical species. (**a**) Feasibility scale index of the coalesced community versus parental competition (uptake-matrix similarity). (**b**) Coalesced-community stability versus parental competition. (**c**) Feasibility scale index versus parental cooperation (effective leakage). (**d**) Coalesced-community stability versus parental cooperation. The overall geometry mirrors Fig. A2, indicating that coalescing structurally heterogeneous parental communities still increases feasibility and stability under 50% overlap. However, the surfaces span a narrower range than in Fig. A2, implying that greater species overlap reduces inter-community heterogeneity and compresses the attainable feasibility/stability landscape, thereby diminishing the effect of coalescence.

### C Traits of the top 10 and bottom 10 species after coalescence

Trait differences between the top 10 and bottom 10 species ranked by abundance in the coalesced community have been further analyzed. As shown in Fig. A4, six traits are examined. Top 10 species exhibit broader resource uptake preferences, and the high variance within this group indicates the presence of multiple broad-spectrum strategies (Fig. A4a). They also display higher and more widely distributed niche overlap (Fig. A4b), suggesting that these species gain abundance primarily through competition rather than by avoiding others. In addition, top 10 species exert stronger effective interactions on the surviving subsystem (Fig. A4d), which in a competitive context reflects stronger suppression and resource depletion effects on other species. Their intrinsic growth rates are markedly higher and cluster near the upper bound, whereas bottom-ranked species show lower and more dispersed values (Fig. A4e). In contrast, bottom 10 species exhibit more specialized resource use (Fig. A4f). They resemble beneficiaries of leaked metabolites rather than primary resource competitors (Fig. A4c). This pattern suggests that in the presence of broad-spectrum competitive species, high specialization alone cannot sustain growth. Cross-feeding functions as a supplementary strategy that maintains low-abundance persistence. In summary, the relatively more competitive parental community contributes dominant species characterized by broad uptake and strong suppressive effects, which occupy most of the biomass and shape stability. The relatively more cooperative parental community contributes a set of specialized, cross-feeding-dependent species that expand functional and species diversity without altering the dominant abundance structure. This result is supported by empirical studies (**???**).

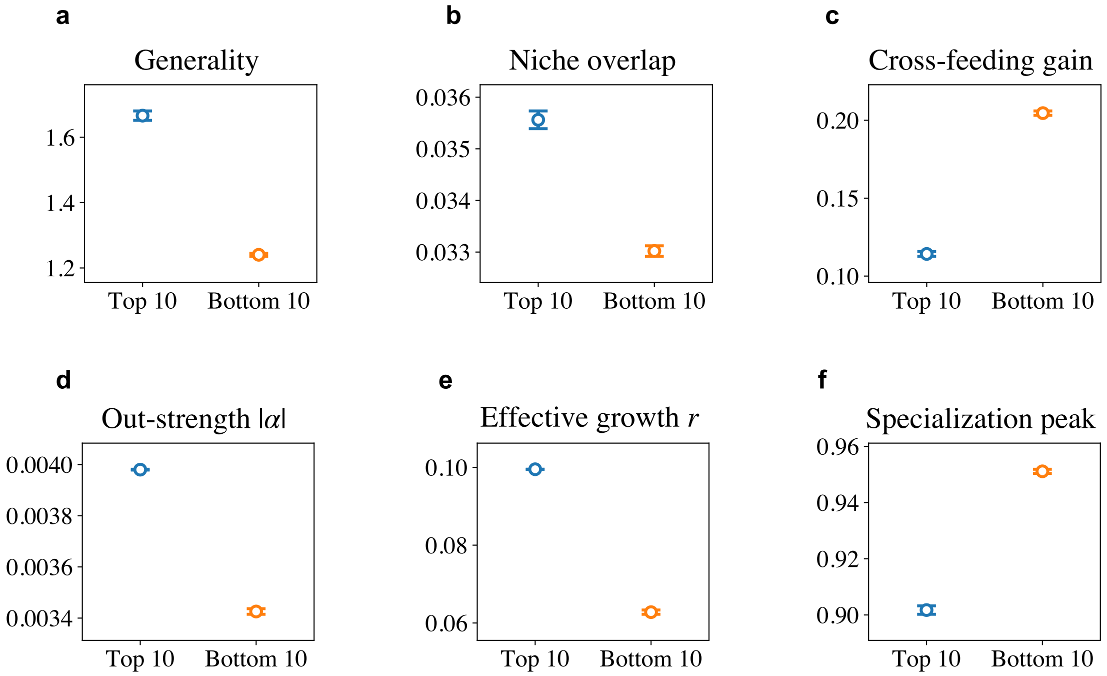


Figure A4: Traits of the top 10 and bottom 10 species by abundance in the coalesced community. For each coalesced community, trait values were first averaged across the top 10 and bottom 10 species, respectively. Points indicate the mean across coalesced communities, and error bars represent 95% confidence intervals of the mean. (**a**) Generality quantifies the breadth of resource use and is calculated from the entropy of the uptake vector; higher values indicate broader resource utilization. (**b**) Niche overlap measures the similarity between a species' uptake pattern and those of other community members. It is computed analogously to the similarity metric in Appendix E; higher values indicate stronger niche overlap. (**c**) Cross-feeding gain reflects the potential benefit a species derives from metabolites leaked by others; higher values indicate greater dependence on cross-feeding. The calculation follows the effective leakage metric described in Appendix E. (**d**) Out-strength $|\alpha|$ is derived from the gLV interaction matrix and measures the total interaction strength a species exerts on other surviving members; higher values indicate a more dominant position in the interaction network. (**e**) Effective growth $r$ represents the net growth tendency of a species under the effective gLV mapping; higher values indicate a greater ability to maintain positive growth. (**f**) Specialization peak describes the concentration of resource preference; higher values indicate stronger specialization on a single resource.

### D Stability analysis

In the MiCRM, the system state is described by the concatenated state vector

$$\mathbf{S} = (\mathbf{C}^\top, \mathbf{R}^\top)^\top, \tag{A1}$$

where $\mathbf{C} \in \mathbb{R}^N$ and $\mathbf{R} \in \mathbb{R}^M$. Linearizing the dynamics around a fixed point $(\hat{\mathbf{C}}, \hat{\mathbf{R}})$ gives

$$\frac{d\Delta\mathbf{S}(t)}{dt} = \mathbf{J}\,\Delta\mathbf{S}(t), \tag{A2}$$

where $\mathbf{J}$ is the $(N+M) \times (N+M)$ Jacobian matrix evaluated at $(\hat{\mathbf{C}}, \hat{\mathbf{R}})$.

In the MiCRM, consumer interactions are mediated explicitly by resource dynamics. Let

$$f_i \equiv \frac{dC_i}{dt}, \qquad g_\alpha \equiv \frac{dR_\alpha}{dt}, \tag{A3}$$

and define the vector fields $\mathbf{f} = (f_1, \ldots, f_N)^\top$ and $\mathbf{g} = (g_1, \ldots, g_M)^\top$. The Jacobian then has the block form

$$\mathbf{J} = \begin{bmatrix} \mathbf{J}_{CC} & \mathbf{J}_{CR} \\ \mathbf{J}_{RC} & \mathbf{J}_{RR} \end{bmatrix} = \begin{bmatrix} \dfrac{\partial \mathbf{f}}{\partial \mathbf{C}} & \dfrac{\partial \mathbf{f}}{\partial \mathbf{R}} \\ \dfrac{\partial \mathbf{g}}{\partial \mathbf{C}} & \dfrac{\partial \mathbf{g}}{\partial \mathbf{R}} \end{bmatrix}. \tag{A4}$$

The four blocks are given below.

**Consumer-consumer block $\mathbf{J}_{CC}$.** Since $f_i$ depends on other consumers only indirectly through resources, the direct partial derivative with respect to $C_j$ is

$$\frac{\partial f_i}{\partial C_j} = \delta_{ij} \left[ \sum_{\alpha=1}^{M} R_\alpha u_{i\alpha}(1-\lambda_i) - m_i \right]. \tag{A5}$$

Evaluating this at the fixed point gives

$$(\mathbf{J}_{CC})_{ij} = \delta_{ij} \left[ \sum_{\alpha=1}^{M} \hat{R}_\alpha u_{i\alpha}(1-\lambda_i) - m_i \right]. \tag{A6}$$

For any surviving species with $\hat{C}_i > 0$, the bracketed term vanishes at equilibrium, and hence the corresponding diagonal entry is zero.

**Consumer-resource block $\mathbf{J}_{CR}$.** This block describes the response of consumer growth to resource variation:

$$\frac{\partial f_i}{\partial R_\gamma} = C_i u_{i\gamma}(1-\lambda_i). \tag{A7}$$

Thus, at the fixed point,

$$(\mathbf{J}_{CR})_{i\gamma} = \hat{C}_i u_{i\gamma}(1-\lambda_i). \tag{A8}$$

**Resource-consumer block $\mathbf{J}_{RC}$.** This block captures both resource depletion by consumption and replenishment through leakage:

$$\frac{\partial g_\alpha}{\partial C_k} = -R_\alpha u_{k\alpha} + \sum_{\beta=1}^{M} R_\beta u_{k\beta} l^k_{\beta\alpha}. \tag{A9}$$

Evaluated at the fixed point,

$$(\mathbf{J}_{RC})_{\alpha k} = -\hat{R}_\alpha u_{k\alpha} + \sum_{\beta=1}^{M} \hat{R}_\beta u_{k\beta} l^k_{\beta\alpha}. \tag{A10}$$

**Resource-resource block $\mathbf{J}_{RR}$.** This block describes the local resource dynamics arising from dilution, consumption, and leakage:

$$\frac{\partial g_\alpha}{\partial R_\gamma} = \sum_{i=1}^{N} C_i u_{i\gamma} l^i_{\gamma\alpha} - \delta_{\alpha\gamma}\left(\omega_\alpha + \sum_{i=1}^{N} C_i u_{i\alpha}\right). \tag{A11}$$

Hence, at the fixed point,

$$(\mathbf{J}_{RR})_{\alpha\gamma} = \sum_{i=1}^{N} \hat{C}_i u_{i\gamma} l^i_{\gamma\alpha} - \delta_{\alpha\gamma}\left(\omega_\alpha + \sum_{i=1}^{N} \hat{C}_i u_{i\alpha}\right). \tag{A12}$$

The fixed point is locally asymptotically stable if all eigenvalues of $\mathbf{J}$ have negative real parts, that is,

$$\max_k \mathrm{Re}(\mu_k) < 0, \tag{A13}$$

where $\mu_k$ denotes the eigenvalues of $\mathbf{J}$.

### E Modularity driven by cooperation-competition balance

Let $\mathbf{U} \in \mathbb{R}^{N\times M}$ denote the uptake matrix generated for a given community, where $N$ is the number of species and $M$ is the number of resources. Each row vector $\mathbf{u}_i \in \mathbb{R}^M$ corresponds to the uptake profile of species $i$. The pairwise cosine similarity between the uptake vectors of species $i$ and $j$ is defined as:

$$\mathrm{cos\_sim}(\mathbf{u}_i, \mathbf{u}_j) = \frac{\mathbf{u}_i \cdot \mathbf{u}_j}{\|\mathbf{u}_i\|\,\|\mathbf{u}_j\|}, \tag{A14}$$

where $\cdot$ denotes the Euclidean inner product and $|\mathbf{u}_i|$ is the Euclidean norm of $\mathbf{u}_i$. The average row-wise similarity across all species is then computed as:

$$\overline{\mathrm{cos\_sim}} = \frac{2}{N(N-1)} \sum_{1\le i<j\le N} \mathrm{cos\_sim}(\mathbf{u}_i, \mathbf{u}_j), \tag{A15}$$

which represents the mean cosine similarity over all distinct species pairs within the community. Higher values indicate more homogeneous uptake strategies among species, whereas lower values reflect greater differentiation in resource utilization. Figure A5a shows the effectiveness of parameter $b$ in similarity adjustment.

The relationship between the $b$ and the effective facilitation strength within the community is further quantified. Let $\mathbf{U} \in \mathbb{R}^{N\times M}$ denote the species-resource uptake matrix, and $\mathbf{L} \in \mathbb{R}^{N\times M\times M}$ represent the species-specific leakage tensors, where $L^i_{\alpha\beta}$ corresponds to the fraction of resource $\alpha$ leaked by species $i$ that contributes to resource $\beta$. For each species $i$, the effective facilitation is computed as:

$$F_i = \sum_{\beta=1}^{M}\left(\sum_{\alpha\neq\beta} L^i_{\alpha\beta} \cdot \frac{1}{N-1}\sum_{j\neq i} u_{j,\beta}\right), \tag{A16}$$

where the diagonal elements of $\mathbf{L}^i$ are excluded to consider only cross-resource transformation. The community-level effective facilitation is then obtained by averaging across all species:

$$\overline{F} = \frac{1}{N}\sum_{i=1}^{N} F_i. \tag{A17}$$

Figure A5b illustrates how parameter $b$ controls effective facilitation.

Uptake vector similarity is a normalized pairwise statistic that is sensitive to discrete modules controlled by $b$, while effective facilitation is a summed cross-feeding flux that averages over many interactions in both $\mathbf{U}$ and $\mathbf{L}$ and therefore appears much closer to linear.

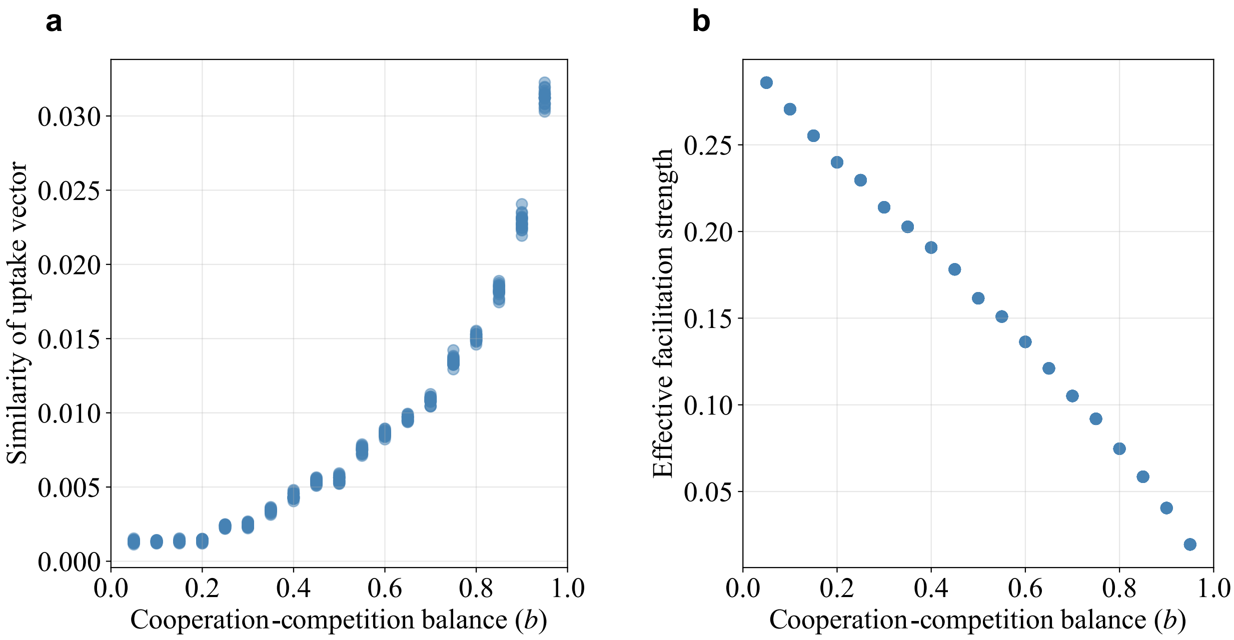


Figure A5: Relationship between the competition–cooperation balance parameter $b$ and uptake vector similarity (**a**) and effective metabolic facilitation (**b**). Across the range of $b$, we compute the mean cosine similarity of uptake vectors in the uptake matrix $\mathbf{U}$ to quantify the similarity of resource preferences among species within a community. In cooperative communities ($b \to 0$), each species exhibits a distinct resource preference, resulting in low uptake similarity. In contrast, in competitive communities ($b \to 1$), species compete more uniformly for all resources, leading to higher uptake similarity. We define effective metabolic facilitation by first removing the diagonal elements of the leakage matrix $\mathbf{L}$, thereby considering only metabolite leakage that benefits other species. The remaining leakage terms are then multiplied by the uptake of other species to quantify the potential for metabolic byproducts to be utilized by community members. The results show that effective metabolic facilitation decreases as communities transition from cooperative to competitive regimes under increasing $b$.

### F Derivation of a gLV Approximation from the MiCRM

We consider a MiCRM governed by

$$\begin{aligned}\frac{dC_i}{dt} &= C_i\left[\sum_{\alpha=1}^{M} R_\alpha\, u_{i\alpha}(1-\lambda_i) - m_i\right],\\ \frac{dR_\alpha}{dt} &= \rho_\alpha - \omega_\alpha R_\alpha - \sum_{i=1}^{N} C_i R_\alpha u_{i\alpha} + \sum_{i=1}^{N}\sum_{\beta=1}^{M} C_i R_\beta u_{i\beta} l^i_{\beta\alpha}.\end{aligned} \tag{A18}$$

Here, for each species $i$ and consumed resource $\beta$, the leakage fractions satisfy

$$\sum_{\alpha=1}^{M} l^i_{\beta\alpha} = \lambda_i, \tag{A19}$$

where $u_{i\alpha}$ denotes the uptake rate of resource $\alpha$ by consumer $i$, $m_i$ is the maintenance cost of consumer $i$, and $\rho_\alpha$ and $\omega_\alpha$ are the external supply and dilution rates of resource $\alpha$, respectively.

Our goal is to derive a local generalized Lotka–Volterra (gLV) approximation around an equilibrium point, of the form

$$\frac{dC_i}{dt} = C_i\left(r_i + \sum_j \alpha_{ij} C_j\right), \tag{A20}$$

and to obtain explicit expressions for the effective intrinsic growth rates $r_i$ and interaction coefficients $\alpha_{ij}$.

#### F.1 Time-Scale Separation and Quasi-Steady-State Approximation

We assume that resource dynamics are much faster than consumer dynamics. Under this quasi-steady-state approximation,

$$\frac{dR_\alpha}{dt} = 0. \tag{A21}$$

Thus, for each resource $\alpha$,

$$\rho_\alpha - \omega_\alpha R_\alpha - \sum_i C_i R_\alpha u_{i\alpha} + \sum_i \sum_\beta C_i R_\beta u_{i\beta} l^i_{\beta\alpha} = 0. \tag{A22}$$

Hence the resource abundance can be expressed as an instantaneous function of the consumer abundances, denoted by

$$R_\alpha = R^*_\alpha(\mathbf{C}). \tag{A23}$$

Substituting this quasi-steady-state relation into the consumer equation gives

$$\frac{dC_i}{dt} = C_i\left[\sum_\alpha u_{i\alpha}(1-\lambda_i) R^*_\alpha(\mathbf{C}) - m_i\right]. \tag{A24}$$

#### F.2 Taylor Expansion Around Equilibrium

Let $(\hat{\mathbf{C}}, \hat{\mathbf{R}})$ be an equilibrium of the MiCRM, with

$$\hat{R}_\alpha = R^*_\alpha(\hat{\mathbf{C}}). \tag{A25}$$

We perform a first-order Taylor expansion of $R^*_\alpha(\mathbf{C})$ around $\mathbf{C} = \hat{\mathbf{C}}$:

$$R^*_\alpha(\mathbf{C}) \approx \hat{R}_\alpha + \sum_j \left.\frac{\partial R^*_\alpha}{\partial C_j}\right|_{\hat{\mathbf{C}}} (C_j - \hat{C}_j). \tag{A26}$$

Substituting this expansion into the consumer dynamics yields

$$\begin{aligned}\frac{dC_i}{dt} &\approx C_i \left\{ \sum_\alpha u_{i\alpha}(1-\lambda_i) \left[ \hat{R}_\alpha + \sum_j \frac{\partial R^*_\alpha}{\partial C_j}\bigg|_{\hat{\mathbf{C}}} (C_j - \hat{C}_j) \right] - m_i \right\} \\ &= C_i \left\{ \underbrace{\left[ \sum_\alpha u_{i\alpha}(1-\lambda_i)\hat{R}_\alpha - m_i \right]}_{r'_i} + \sum_j \left[ \sum_\alpha u_{i\alpha}(1-\lambda_i) \frac{\partial R^*_\alpha}{\partial C_j}\bigg|_{\hat{\mathbf{C}}} \right] (C_j - \hat{C}_j) \right\}.\end{aligned} \tag{A27}$$

We therefore define the effective interaction coefficient as

$$\alpha_{ij} \equiv \sum_\alpha u_{i\alpha}(1-\lambda_i) \frac{\partial R^*_\alpha}{\partial C_j}\bigg|_{\hat{\mathbf{C}}}. \tag{A28}$$

Collecting the constant terms then gives the effective intrinsic growth rate

$$r_i = \sum_\alpha u_{i\alpha}(1-\lambda_i)\hat{R}_\alpha - m_i - \sum_j \alpha_{ij}\hat{C}_j. \tag{A29}$$

#### F.3 Calculation of the Resource Sensitivity $\partial R^*_\alpha / \partial C_j$

To compute the partial derivatives, we define the resource nullcline

$$F_\alpha(\mathbf{C}, \mathbf{R}) \equiv \rho_\alpha - \omega_\alpha R_\alpha - \sum_i C_i R_\alpha u_{i\alpha} + \sum_i \sum_\beta C_i R_\beta u_{i\beta} l^i_{\beta\alpha}. \tag{A30}$$

At quasi-steady state, $F_\alpha(\mathbf{C}, \mathbf{R}^*(\mathbf{C})) = 0$. By the implicit function theorem,

$$\frac{\partial F_\alpha}{\partial C_j} + \sum_\gamma \frac{\partial F_\alpha}{\partial R_\gamma} \frac{\partial R^*_\gamma}{\partial C_j}\bigg|_{\hat{\mathbf{C}}} = 0, \tag{A31}$$

evaluated at $(\mathbf{C}, \mathbf{R}) = (\hat{\mathbf{C}}, \hat{\mathbf{R}})$.

Differentiating $F_\alpha$ with respect to $C_j$ gives

$$\frac{\partial F_\alpha}{\partial C_j}\bigg|_{(\hat{\mathbf{C}}, \hat{\mathbf{R}})} = -\hat{R}_\alpha u_{j\alpha} + \sum_\beta \hat{R}_\beta u_{j\beta} l^j_{\beta\alpha}. \tag{A32}$$

Next, the Jacobian with respect to the resource variables is

$$D_{\alpha\gamma} \equiv \frac{\partial F_\alpha}{\partial R_\gamma}\bigg|_{(\hat{\mathbf{C}}, \hat{\mathbf{R}})} = -\delta_{\alpha\gamma}\left( \omega_\alpha + \sum_i \hat{C}_i u_{i\alpha} \right) + \sum_i \hat{C}_i u_{i\gamma} l^i_{\gamma\alpha}. \tag{A33}$$

Therefore,

$$\sum_\gamma D_{\alpha\gamma} \frac{\partial R^*_\gamma}{\partial C_j}\bigg|_{\hat{\mathbf{C}}} = \hat{R}_\alpha u_{j\alpha} - \sum_\beta \hat{R}_\beta u_{j\beta} l^j_{\beta\alpha}. \tag{A34}$$

After matrix inversion,

$$\frac{\partial R^*_\gamma}{\partial C_j}\bigg|_{\hat{\mathbf{C}}} = \sum_\alpha D^{-1}_{\gamma\alpha} \left[ \hat{R}_\alpha u_{j\alpha} - \sum_\beta \hat{R}_\beta u_{j\beta} l^j_{\beta\alpha} \right]. \tag{A35}$$

##### F.4 Final Expressions

Substituting this result into the definition of $\alpha_{ij}$, we obtain

$$\alpha_{ij} = \sum_{\alpha} u_{i\alpha}(1-\lambda_i) \left\{ \sum_{\gamma} D^{-1}_{\alpha\gamma} \left[ \hat{R}_{\gamma} u_{j\gamma} - \sum_{\beta} \hat{R}_{\beta} u_{j\beta} l^{j}_{\beta\gamma} \right] \right\}. \tag{A36}$$

Similarly, the effective intrinsic growth rate is

$$r_i = \sum_{\alpha} u_{i\alpha}(1-\lambda_i)\hat{R}_{\alpha} - m_i - \sum_{j} \alpha_{ij}\hat{C}_j. \tag{A37}$$
